# scEMB: Learning context representation of genes based on large-scale single-cell transcriptomics

**DOI:** 10.1101/2024.09.24.614685

**Authors:** Kang-Lin Hsieh, Yan Chu, Xiaoyang Li, Patrick G. Pilié, Yulin Dai

## Abstract

**Background:** The rapid advancement of single-cell transcriptomic technologies has led to the curation of millions of cellular profiles, providing unprecedented insights into cellular heterogeneity across various tissues and developmental stages. This growing wealth of data presents an opportunity to uncover complex gene-gene relationships, yet also poses significant computational challenges.

**Results:** We present scEMB, a transformer-based deep learning model developed to capture context-aware gene embeddings from large-scale single-cell transcriptomics data. Trained on over 30 million single-cell transcriptomes, scEMB utilizes an innovative binning strategy that integrates data across multiple platforms, effectively preserving both gene expression hierarchies and cell-type specificity. In downstream tasks such as batch integration, clustering, and cell type annotation, scEMB demonstrates superior performance compared to existing models like scGPT and Geneformer. Notably, scEMB excels *in silico* correlation analysis, accurately predicting gene perturbation effects in CRISPR-edited datasets and microglia state transition, identifying a few known Alzheimer’s disease (AD) risks genes in top gene list. Additionally, scEMB offers robust fine-tuning capabilities for domain-specific applications, making it a versatile tool for tackling diverse biological problems such as therapeutic target discovery and disease modeling.

**Conclusions:** scEMB represents a powerful tool for extracting biologically meaningful insights from complex gene expression data. Its ability to model *in silico* perturbation effects and conduct correlation analyses in the embedding space highlights its potential to accelerate discoveries in precision medicine and therapeutic development.

## INTRODUCTION

The rapid advancement of single-cell technologies has enabled international consortia, including the Human Cell Atlas (HCA)^1^, Tabula Sapiens^2^, Human Cell Landscape^3^, and the more recent 95 million CELLxGENE corpus^4^, to characterize the cellular heterogeneity in major human tissues and organ systems in different developmental stages. However, managing and extracting meaning and value from such large-scale data is highly challenging and this technical gap creates opportunities for researchers to innovate and develop advanced computational methods capable of harnessing the full potential of these vast datasets, enabling deeper biological understanding and practical applications.

Clustered regularly interspaced short palindromic repeats (CRISPR) system is a genetic engineering tool for gene editing that can activate (CRISPRa), repress (CRISPRi), or modify gene expressions for specific genes^5^. This technology allows us to understand the systematic effects of perturbing one or multiple genes in different cell types, which provides ideal data to learn the relationship between different perturbations^6,7^. However, genome-scale gene perturbations and their potential combinations have been impeded by the cost, emphasizing the need for novel cost-efficient methods to prioritize their effects.

Recent advances in artificial intelligence (AI), such as variational autoencoder (VAE) models^8–10^, which explicitly learn low-dimensional embeddings from single-cell transcriptome to capture the biological meaning embedding. Moreover, these embedding-based methods^11^ [Autoencoder^12^, VAE^13–16^, GNN^17^], has been widely used to along with the cellular perturbation tasks along with single-cell CRISPR data, to compress the transcriptomics data (mainly single-cell) into the latent space and compare the effects of different perturbations and possess a more accurate molecular measurement as well as in silico-induced perturbations^18,19^. However, these methods^18,19^ were mainly task-specific and the perturbation effect were encoded on different embedding systems^9^, hindering the transfer learning performance.

More recently, large-language models (LLM) such as GPT^20,21^, have revolutionized various fields by leveraging deep neural networks trained on vast text datasets. These models generalize knowledge from pretraining, enabling efficient task transfer with minimal data. The self-attention mechanism enhances their ability to focus on relevant information, improving predictions across diverse applications.

In single-cell data analysis, transformer-based models offer a promising approach to overcoming batch effects through batch-unaware pretraining, which has proven resilient to some technical artifacts. For instance, models like Universal Cell Embeddings (UCE)^22^ and GeneCompass^23^ have integrated molecular profiles across studies, tissues, and species, enabling cell type annotations to transfer across previously unseen species. More recent Geneformer^25^, scGPT^24^,, and scFoundation^24^ trained the foundation model from large-scale single-cell corpus and offered transfer learning features^25^ to makes the transformer an ideal backbone for inferring unseen perturbations^17,26,27^, showing great promise to mitigate previous limitations.

Despite significant advances, a key gap remains in applying transformer models specifically to single-cell data. While transformers have shown promise in handling large-scale transcriptomic data, optimization for single-cell applications is still needed. To address this, we aim to develop the scEMB model to explore broader application scenarios in single-cell foundation models and establish best practices across diverse datasets. Additionally, we will investigate the potential of in silico perturbation prediction using both zero-shot and fine-tuned models, enabling more accurate biological insights and expanding applications in disease modeling and therapeutic discovery.

## RESULT

### Overview of scEMB

scEMB is an attention-based deep learning model pretrained on large-scale transcriptomic data to capture complex biological networks. It uses self-attention to focus on the most critical genes expressed in each single cell, optimizing predictive accuracy through various learning objectives. scEMB is designed to capture complex biologically meaning alteration underlying cell type state transition and perturbation response (Fig. 1).

**Fig. 1.**
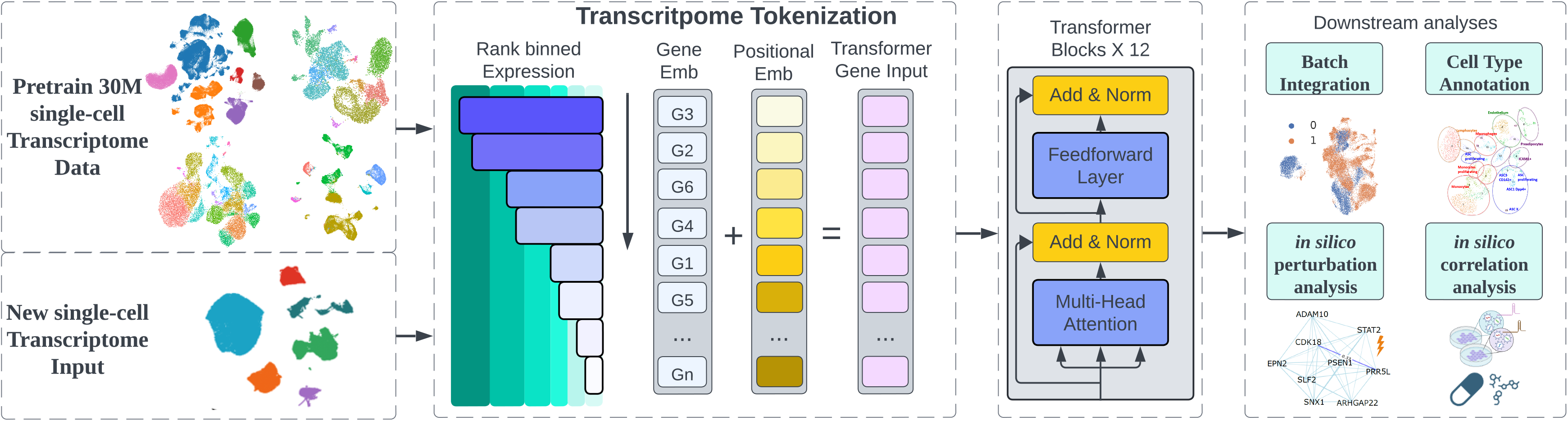
Framework of scEMB. The 30M single-cell transcriptome dataset, curated by CZ CELLxGENE Discover, was processed by normalizing gene expression to counts per million and applying a log1p transformation. We employed a binned expression strategy adapted from scGPT, binning genes into 100 intervals and ranking them based on their real expression values. Leveraging a BERT model as the backbone, we engineered the positional embeddings to preserve the order of real expression values. The resulting concatenated embeddings were fed into 12 transformer blocks, training the model to capture gene order and generate a gene-attention map to represent cells. During inference, the gene expression data for new cells were tokenized and inputted into the pretrained model to obtain both gene and cell embeddings. These embeddings were then used in various downstream tasks to streamline conventional single-cell analyses. This figure was created using materials adapted from Biorender.com.

scEMB represents each single cell’s transcriptome as a rank-binned gene expression encoding, ranking genes by their expression within individual cells. While this rank-based approach has limitations, such as not fully utilizing precise gene expression measurements from transcript counts, it provides a non-parametric representation of each cell’s transcriptome. This method leverages the vast observations of gene expression across the cell-x-gene 30M dataset to highlight genes that characterize cell states by composing cellular biological networks.

The rank value encoding of each single cell’s transcriptome then passes through twelve transformer encoder units, each consisting of a self-attention layer and a feed-forward neural network layer. Pretraining utilized a masked learning objective, a technique proven in other information domains to enhance the generalizability of foundational knowledge acquired during pretraining, for a broad spectrum of downstream fine-tuning tasks such as batch integration, cell type annotation, *in silico* perturbation, and correlation analysis.

### scEMB shows comparable performance in clustering and batch integration

As the standard tasking in single-cell foundation model, we tested the performance in single-cell clustering using scEMB, scGPT, Geneformer, and unintegrated methods on PBMC 10k dataset^28^. Fig. 2 shows the PBMC 10k dataset before and after dimensionality reduction using UMAP (Uniform Manifold Approximation and Projection), with the legend indicating various cell types, including B cells, CD4 T cells, CD8 T cells, CD14+ Monocytes, Dendritic Cells, FCGR3A+ Monocytes, Megakaryocytes, NK cells, and other. For batch integration and cell type clustering tasks, we present UMAP plots that illustrate cell clustering across different datasets using four methods, with distinct colors indicating various cell types. Below the UMAP plots, a performance comparison table presents various metrics for each method, including isolation score, kBET score, and batch mixing entropy. The table also includes aggregate scores for metrics such as batch correction, biological conservation, and overall performance. This comprehensive visualization enables a direct comparison of how well each method integrates data across batches while preserving biological information.

**Fig. 2.**
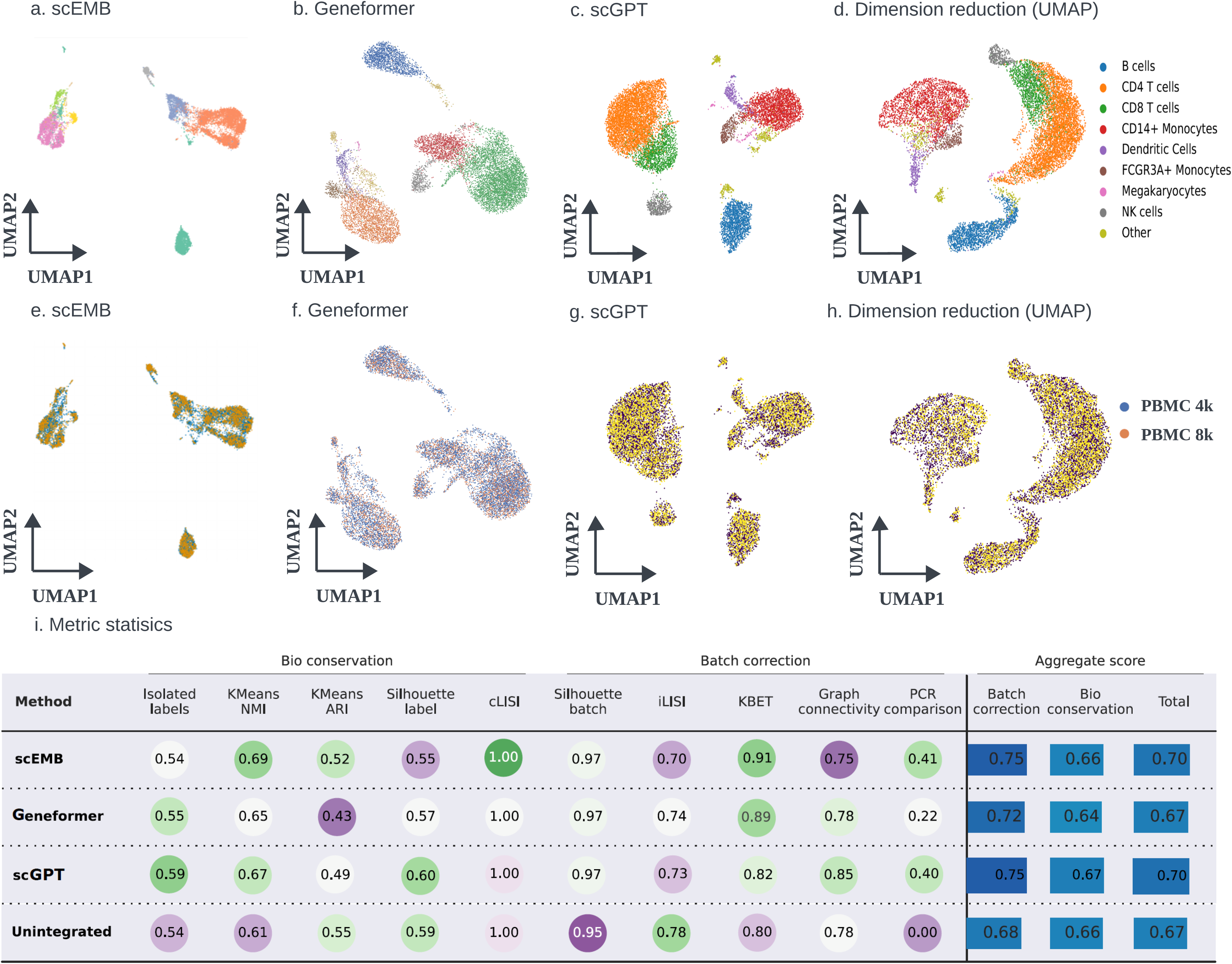
Clustering performance of PBMC10k data. a-d, UMAP plots illustrating clustering performance for scEMB, Geneformer, scGPT, and conventional dimensionality reduction methods on the PBMC10k dataset. e-h, UMAP plots illustrating batch integration performance for scEMB, Geneformer, scGPT, and conventional dimensionality reduction methods on the PBMC10k dataset. i, Five clustering metrics are compared across the four benchmark methods, with scores ranging from 0 to 1, where higher values indicate better performance.

### High consistent performance in cell type annotation task

One of the main advantages of foundation models is their ability to leverage pretraining on large-scale general datasets, enabling fine-tuning for a wide range of downstream tasks, even when the available task-specific data is sparse to generate meaningful predictions. We evaluated scEMB’s performance on cell-type annotation tasks by training a supervised learning model on cell-type annotations from a reference dataset, then predicting cell types in an independent, unseen dataset using two batches of data from MTG brain region dataset^29^.

After inputting the transcriptome into scEMB Encoder, the cell-level embedding generated by scEMB, representing a specific cellular state, can be used to infer cell type. The cell type annotation was trained in one batch of the brain MTG dataset, and test on the other batch (Fig. 3a). To evaluate cell type prediction accuracy, we used confusion matrices for scEMB, scGPT, and Geneformer (Fig. 3b-3d). While all methods performed well, we observed slight differences, especially for less common cell types. This visualization offers both qualitative (UMAP) and quantitative (confusion matrix) insights into scEMB’s effectiveness in predicting cell types from single-cell RNA sequencing data. Overall, all foundation models demonstrated strong potential in accurately annotating cell types using well-annotated reference data, a critical step in single-cell data analysis.

**Fig. 3.**
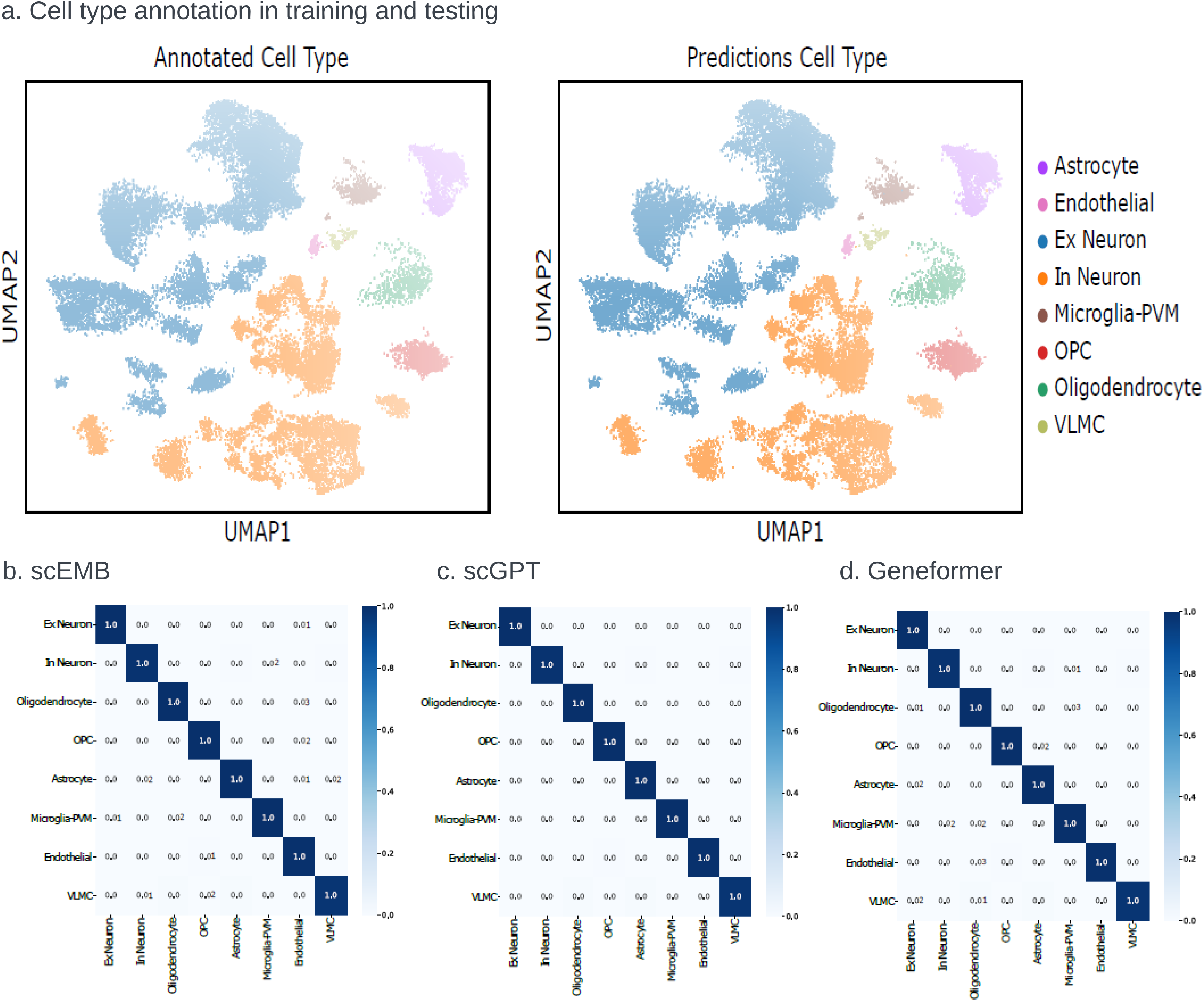
Cell type annotation performance after fine-tuning on brain MTG datasets. We adapted two samples from SEA-AD cohort. a. One sample was used for training the cell type classifier, while the other sample was reserved for testing the classification performance. b-d. We benchmarked the cell type annotation performance of scEMB against scGPT and Geneformer, and presented the results using confusion matrix. Darker blue along the diagonal indicates higher accuracy.

### scEMB *in silico* perturbation analysis identified genes highly consistent with CRISPRi data

As demonstrated by Geneformer and scGPT, single-cell foundation models show great potential in predicting cellular gene expression responses to specific gene perturbations. This highlights a major advantage of transfer learning, which leverages biological knowledge from millions of human cells to improve accuracy and scalability. To investigate potential deleterious effects in response to cellular perturbation, we designed an *in silico* perturbation response prediction task, which was validated using a single-cell CRISPRi dataset with known perturbation outcomes (Fig. 4a). Specifically, we measured these effects using a cosine similarity score calculated from the scEMB-encoded embeddings in an iPSC-derived microglia dataset^30^. Among the 39 perturbed gene conditions (Fig. 4b), we identified the top 10 most accurately predicted *in silico* perturbed genes, which were validated against ground truth CRISPRi data. Among them, we identified a few well know microglia function genes, such CSF1R (Colony Stimulating Factor 1 Receptor), which is critical for the survival, differentiation, and function of microglia^31^. TGFBR1/2 (Transforming Growth Factor Beta Receptor 1/2) TGFBR1/2 are involved in the TGF-β signaling pathway, which is essential for microglial homeostasis and modulation of their inflammatory response. TGF-β helps maintain the quiescent state of microglia in the healthy brain but also plays a role in activating microglia during injury or neurodegenerative processes^32^. Interestingly, the housekeeping gene like AARS (Alanyl-tRNA Synthetase)^33^ demonstrated the highest cosine similarity in predictions, indicating that scEMB may perform well in predicting housekeeping genes. However, this result also suggests that the universal expression of housekeeping genes across cells could be more complex, as highlighted by findings from the single-cell foundation model^27^.

**Fig. 4.**
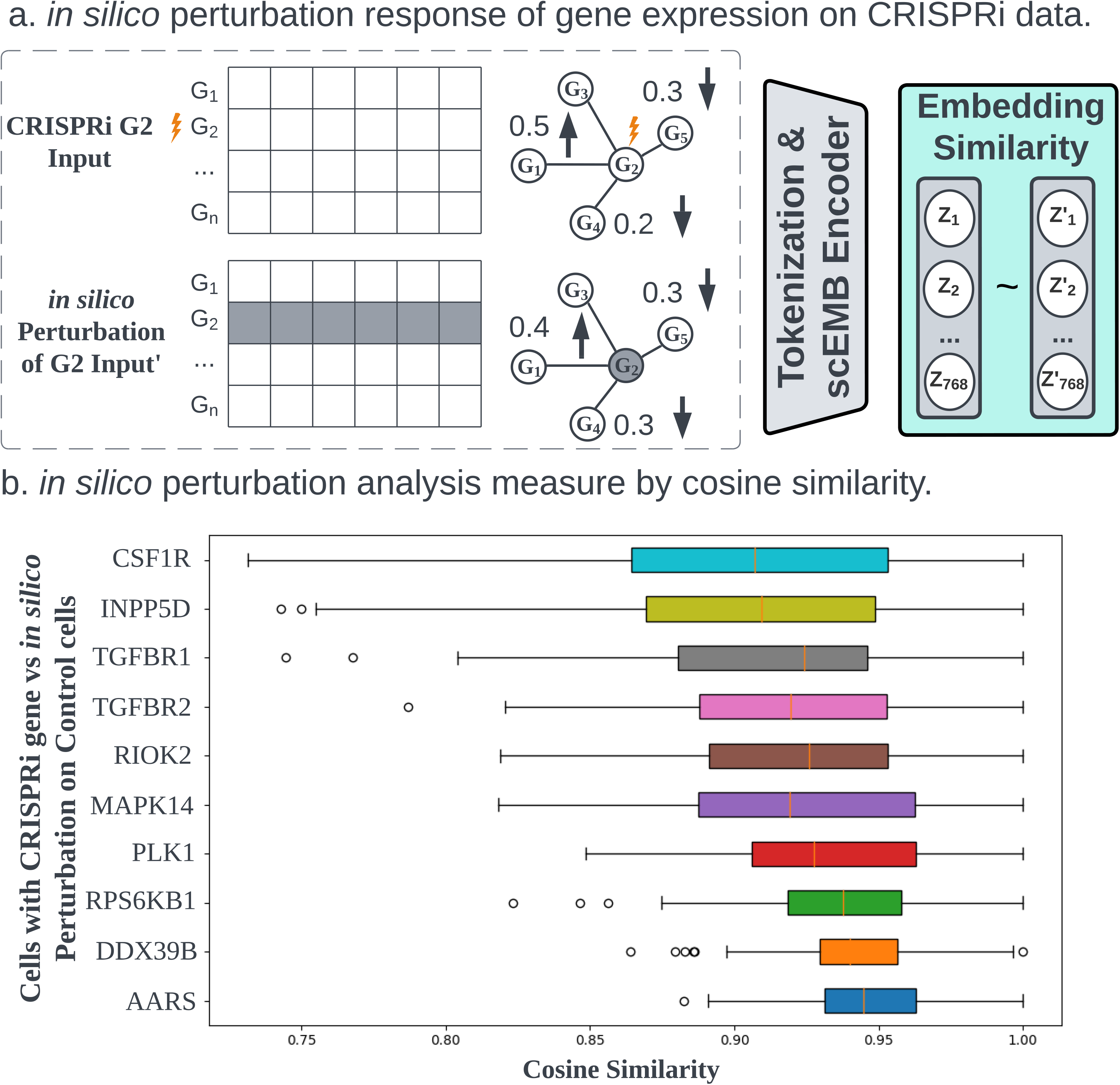
scEMB *in silico* perturbation analysis. a. Diagram of the scEMB *in silico* perturbation analysis framework, from input data processing to the comparison of cellular embedding similarities. To evaluate the performance of the *in silico* perturbation, we used a CRISPRi dataset as the ground truth and measured prediction accuracy by calculating the cosine similarity of the cell embeddings. b. Bar plot showing the cosine similarity of cell embeddings for 10 CRISPR-edited genes, comparing their *in silico* perturbations on controls with the corresponding CRISPRi gene perturbated cells.

### scEMB *in silico* correlation analysis highlighted AD risk genes that might contribute to the AD pathogenesis

As there is still limited real world perturbation data existing, not to mention most perturbation response data or conditions is not included in training in current foundation models^24,27,34^. Therefore, perturbation response predicted from zero-shot learning still requires careful use. We designed a finetune task based alternative approach as revealed in Fig. 5. scEMB *in silico* corrrelation analysis could be applied to identify potential reversed relationship at cellular embedding level between cellular state alteration effect and perturbation effect among single-cell transcriptome data. Here, we first project the cellular state alteration embeddings using scEMB Encode, focusing on microglia from individuals with Alzheimer’s Disease (AD) and cognitively normal (CN) individuals from the Religious Orders Study/Memory and Aging Project (ROS/MAP) cohort^36^. Then, we followed our previously perturbation strategy^10,17^ to integrate pretrained GO Gene GNN node embeddings, allowing us to propagate the impact of perturbations from drug treatment or iPSC-derived CRISPRi data^30^. As illustrated in Fig. 5b, the violin plot presents the top 15 absolute cosine similarity scores, highlighting cellular state alterations in microglia by comparing cells from individuals with Alzheimer’s Disease (AD) to those from cognitively normal controls. Additionally, the plot reveals the effects of gene perturbations on iPSC-derived microglia. Notably, several well-known AD risk genes from genome-wide association study^36^, including PLCG2, SORL1, and TREM2, emerged among the top genes. These findings suggest that the scEMB *in silico* correlation analysis may offer valuable insights into genes involved in disease pathogenesis, providing clues about underlying molecular mechanisms. Furthermore, it points to potential therapeutic targets, as the identified genes could play a pivotal role in modulating microglial function and influencing disease progression in AD.

**Fig. 5.**
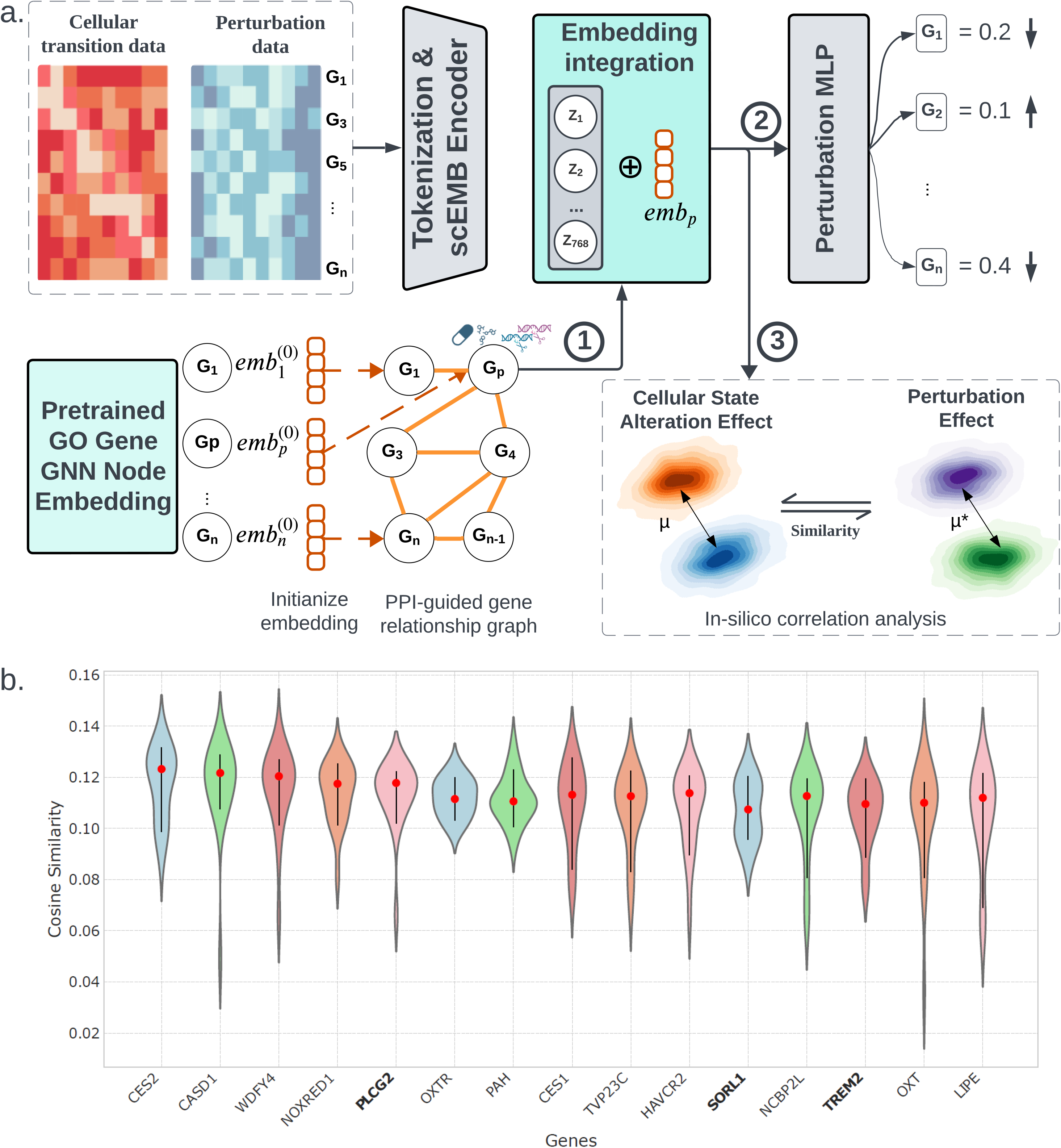
scEMB perturbation response and *in silico* correlation analysis. a. All the single-cell transcriptome were first tokenized and went through scEMB Encoder. The 786-dimensional cellular embedding will be concatenated with gene perturbation embeddings (emb_p_), which are derived from a PPI-guided gene relationship graph (step 1) following our previous method^10^. This graph is built using genetic perturbation data and propagates the impact of perturbations (P on gene G ) across gene-gene relationships informed by gene ontology (GO). scEMB provides two downstream tasks to analyze perturbation effects at both the gene and cellular levels, respectively. For gene-level analysis in step 2, the concatenated embeddings are processed through a two-layer Multilayer Perceptron (MLP), designed to link cell embeddings to gene features. The model outputs altered gene expression profiles, representing the predicted overall impact of perturbations on other genes. For cellular level *in silico* correlation analysis, we designed to test the correlation analysis between celllular state alteration effect and Perturbation effect, and therefore capture to a potenial reversing effect might provide the insight of theraputic targets (step 3). b. The violin plot shows the top 15 absolute cosine similarity scores for cellular state alterations in microglia, comparing cells from Alzheimer’s Disease (AD) with cells from controls, as well as the effects of gene perturbations on iPSC-derived microglia. The absolute cosine similarity measures the similarity between these two effects in 768 dimensions. The x- axis represents the perturbed genes, with AD risk genes highlighted in bold. This figure was created using materials adapted from Biorender.com.

## METHODS AND MATERIALS

### scEMB architecture

scEMB comprises twelve transformer encoder units, each containing a self-attention layer and a feed-forward neural network layer. The model’s key parameters include an input size of 2,048, embedding dimensions of 768, 12 attention heads, and a feed-forward network size of 3,072. The input size was maximized to capture the most context possible through full attention, based on the typical number of genes detected in each cell in the pretraining dataset. To speed up the training process for this large dataset, scEMB employed Scaled Dot-Product Attention (SDPA) across the entire 2,048 input size. The model’s depth was determined by the maximum level for which sufficient pretraining data was available. Other minor parameters include the Gaussian Error Linear Unit as the activation function, a dropout probability of 0.1 for fully connected layers, and a dropout ratio of 0.1 for attention probabilities. The weight matrices were initialized with a standard deviation of 0.02, and an epsilon value of 1 × 10⁻¹² was used for layer normalization. The model was built in PyTorch and utilized the Huggingface Transformers library for configuration, data loading, and training.

### scEMB pretraining

The primary pretraining objective of scEMB is masked language modeling. This approach, proven effective in various fields, enhances the generalizability of foundational knowledge acquired during pretraining, benefiting a wide range of downstream fine-tuning objectives.

During pretraining, 15% of the genes within each transcriptome are masked. The model then learns to predict which gene should occupy each masked position in that specific cell state, using the context provided by the remaining unmasked genes. This self-supervised approach can be accomplished on completely unlabeled data, allowing the inclusion of large amounts of training data without being restricted to samples with accompanying labels.

The pretraining process utilized optimized hyperparameters to enhance model performance. These included a maximum learning rate of 1 × 10⁻L, a linear learning scheduler with warmup, and the Adam optimizer with weight decay fix. Additionally, 10,000 warmup steps were employed, along with a weight decay of 0.001 and a batch size of 8. These carefully selected parameters contributed to the model’s effective training. During model training, we adapted a custom tokenizer from the Huggingface Transformers library to implement dynamic, length-grouped padding. This approach minimized computation on padding and achieved a 29.4× speedup in pretraining. The process involves randomly sampling a megabatch, then ordering minibatches by their length in descending order. These minibatches are dynamically padded, reducing wasted computation on padding due to their grouped lengths. The distributed training is implemented by Deepspeed, which partitions parameters, gradients, and optimizer states across available GPUs. Overall, pretraining was achieved in approximately 10 days, distributed across one node with 8 Nvidia H100 96GB GPUs.

### scEMB finetuning

Fine-tuning scEMB involves initializing the model with pretrained scEMB weights and adding a final task-specific transformer layer. This process aims to recalibrate the distribution between the pre-trained model and the task-specific dataset. The recalibration can be divided into two types: 1) using a small dataset to adjust the layers within the foundation model, or 2) fine-tuning external knowledge or embeddings, generated either from a task-specific dataset or expert knowledge, to feed into the foundation model. In the first type of fine-tuning, the number of frozen layers must be carefully considered. Based on our experience and suggestions from relevant literature, applications more closely related to the pretraining objective benefit from freezing more layers. This prevents overfitting to limited task-specific data. Conversely, applications that diverge more from the pretraining objective benefit from fine-tuning more layers, optimizing performance for the new task. In the second type of fine-tuning, the extraction of features and appropriate use of expert knowledge still need comprehensive discussion. The task-specific dataset underwent the same preprocessing pipeline as scEMB. To demonstrate the effectiveness of the pretrained scEMB model in enhancing the predictive performance of downstream fine-tuning tasks, we employed consistent fine-tuning hyperparameters across all applications. Special attention must be given to determining the appropriate number of frozen layers during the fine-tuning process.

Several applications of scEMB have been demonstrated, including cell type annotations, batch corrections, and perturbation effects. These applications further test the two types of recalibration. For cell type annotations and batch corrections, we used the PBMC dataset. In both subtasks, we tested LoRA (Low-Rank Adaptation) for the first type of fine-tuning and Prefix Tuning for the second type. The key difference is that LoRA aligns the model with a new dataset, while Prefix Tuning aligns the external knowledge vector, typically called an embedding, with a new dataset. Consequently, the cell representation from LoRA fine-tuning uses cell gene expression, whereas Prefix Tuning combines external knowledge and cell gene expression. The cell representation is the average of hidden layer outputs for all 12 layers. The benchmarks for cell type annotation were AUROC, F1 score, and accuracy. For batch corrections, we used the following metrics: Isolated labels, KMeans NMI, KMeans ARI, Silhouette label, cLISI, Silhouette batch, iLISI, KBET, Graph connectivity, and PCR comparison.

### *in silico* perturbation analysis

To evaluate the model’s ability to capture gene perturbation effects, we conducted an *in silico* perturbation analysis, focusing on cosine similarity between cell embeddings of *in silico* perturbed control cells and CRISPR-edited cells. Cell embeddings, representing each cell as a 768-dimensional vector, were obtained by averaging the encoder output. For control cells, we masked the CRISPR-targeted gene, then calculated the cosine similarity between control and perturbed groups. A high cosine similarity suggests a minimal effect on the gene, indicating less significant perturbation. Conversely, lower similarity highlights more impactful gene changes. This evaluation method provides insight into the genes most significantly affected by CRISPR perturbations. The results, as shown in Fig. 4, demonstrate how cosine similarity can effectively reflect gene impact, where high similarity indicates low gene perturbation impact and vice versa. This approach showcases the model’s precision in detecting subtle yet significant gene expression changes after perturbations.

### *in silico* correlation analysis

To further investigate the model’s capability to recognize the similarity between disease state transitions and gene perturbation effects, we conducted an *in silico* correlation analysis. Single-cell transcriptomes from non-targeting controls (NTC) in the Alzheimer’s Disease (AD) dataset were tokenized and processed through the scEMB encoder, producing 768-dimensional cell embeddings. These embeddings were combined with gene perturbation embeddings (emb_p_), which were derived from a protein-protein interaction (PPI)-guided gene relationship graph, informed by gene ontology (GO). The graph utilized a 3-layer graph attention network (GAT) to conduct self-supervised learning, enabling the extraction of embeddings for each gene^10,17^. These combined embeddings were then passed through a two-layer Multilayer Perceptron (MLP), which linked the cell embeddings to gene features, allowing the model to predict altered gene expression profiles. Additionally, the *in silico* correlation analysis was employed to assess the relationship between cellular state alterations and gene perturbation effects. This analysis aimed to identify potential therapeutic targets by capturing the similarity or reversed relationships between disease-related cellular changes and the impact of specific gene perturbations. To evaluate these relationships, we performed a cosine similarity analysis, quantifying the similarity between control and AD cellular states and the effects of gene perturbations on induced pluripotent stem cell (iPSC)-derived microglia. The results, presented in Fig. 5, highlight the AD risk genes, allowing for a better understanding of how perturbations influence the disease state at the cellular level. This approach offers a robust framework for studying perturbation effects and identifying key genes involved in disease mechanisms.

### Benchmark with other models

To comprehensively evaluate the performance of scEMB, we compared its clustering and batch integration capabilities against two other single-cell foundation models, Geneformer^27^ and scGPT^34^, as well as the standard single-cell data analysis approach in Scanpy^37^. These comparisons were conducted in a zero-shot setting using the PBMC 10k dataset^28^ with default configurations. Additionally, we assessed cell type annotation performance in a fine-tuning scenario by conducting a classification task on a brain microglia dataset^36^, comparing scEMB to Geneformer and scGPT.

## Data preprocess

### 30M single-cell transcriptome corpus curation

For pretraining the whole-human foundation model, we sourced data through the Census API from the CELLxGENE portal^38^ (https://cellxgene.cziscience.com/), which provides regularly updated datasets (accessed on May 9, 2024). We included both scRNA-seq and snRNA-seq sequencing protocols, focusing on samples from healthy conditions. To ensure data quality, we filtered out cells expressing fewer than 200 genes or mitochondria gene expression percent > 10 using python library Scanpy^37^. After applying these filters, the final dataset comprised sequencing data from 30 million cells.

### PBMC 10k dataset

The PBMC 10k dataset consists of two single-cell RNA sequencing batches of peripheral blood mononuclear cells (PBMCs) obtained from a healthy donor. This dataset was reanalyzed by Gayoso et al.^28^, identifying 3,346 differentially expressed genes. The first batch consists of 7,982 cells, and the second batch contains 4,008 cells. Cell type annotations were conducted using the R package Seurat^39^ , categorized the cells into nine distinct groups. The preprocessed data was adopted from scGPT^34^ (https://github.com/bowang-lab/scGPT, accessed on June 16, 2024).

### MTG Brain dataset

We incorporated two brain samples from the middle temporal gyrus (MTG) region, provided by the Seattle Alzheimer’s Disease Brain Cell Atlas (SEA-AD) consortium via Amazon AWS Bucket (accessed on August 9, 2024). These two samples (H19.33.004 and H19.30.001) were profiled from two different batches. Cell type annotations were derived from the original study^29^.

### Microglia from ROS/MAP snRNA-seq and polygenic risk score process

We used snRNA-seq data from the Synapse portal (syn2580853, accessed on April 15, 2023), which includes 454 participants from the Religious Orders Study/Memory and Aging Project (ROS/MAP) cohort^36^. Matched whole-genome sequencing (WGS) data were obtained from Synapse (syn11724057, accessed on November 10, 2022). Individual genetic risk was estimated using LDPred2^40^, based on variant effect sizes from Wightman et al.’s GWAS summary statistics^41^. This provided a dataset of 407 individuals with both snRNA-seq and WGS data. To focus on individuals with high polygenic risk scores (PRS) for Alzheimer’s disease, we selected the top 20% PRS, yielding 53 high-risk AD cases and 15 high-risk cognitively normal individuals^42^. Data was processed using R package Seurat^39^ with filters applied for “percent.mt <= 50” and “nFeature_RNA > 200,” and cell type annotations were adopted from the original study.

### CRISPRi iPSC-derived microglia dataset

The CRISPR iPSC-derived microglia dataset was downloaded from GSE178317^30^ (accessed on March 10, 2023) and processed using the CROP-seq pipeline^43^. The gene feature matrix was obtained by alignment and gene expression quantification for scRNA-seq libraries and sgRNA-enriched libraries using Cell Ranger and STAR^44^. 2) The sgRNA matrix was assigned based on the demuxEM algorithm^45^ and a modified z-score cut-off method^46^. We categorized the cells into two groups: those with a single sgRNA and those with two sgRNAs. In total, we identified 39 single sgRNA-targeted genes, each captured by at least 200 cells.

## DISCUSSION

In this study, we introduce scEMB, a novel single-cell transcriptome foundation model developed to extract complex gene-gene interaction information from a large corpus of 30 million human single-cell transcriptomes. A key feature of scEMB is its specially designed tokenization mechanism, which was engineered to generate gene embeddings that are both stable and sensitive. This design allows scEMB to be highly effective across a broad spectrum of downstream tasks, from gene interaction analysis to large-scale biological modeling.

scEMB’s performance has been rigorously evaluated in tasks such as clustering and batch integration, and it demonstrated competitive zero-shot performance across multiple datasets. Remarkably, its performance was on par with leading foundation models like Geneformer and scGPT, reinforcing scEMB’s utility as a powerful tool for a variety of biological applications.

One of the novel contributions of this work is the exploration of scEMB’s potential in perturbation response prediction, using an Alzheimer’s Disease (AD) dataset as a case study. We estimated the cosine similarity between perturbation effects and cellular state transitions, demonstrating how scEMB can model complex cellular responses to external stimuli or genetic modifications. This ability to predict perturbation effects holds significant promise for the study of disease mechanisms and therapeutic interventions.

In our evaluation of fine-tuning approaches, we identified a critical gap in current practices: the lack of a standardized method for fine-tuning large models in biological contexts, particularly in classification tasks. Existing classification tasks often fail to capture the complexity of biological research needs, limiting the practical utility of these models. To address this challenge, we introduced alternative fine-tuning strategies, including Prefix-tuning and LoRA (Low-Rank Adaptation), which are specifically tailored for downstream applications such as cell-type annotation and *in silico* perturbation analysis. These methods offer greater flexibility and more accurate biological insights, especially in tasks that involve nuanced gene expression changes across different conditions.

Furthermore, based on observations from models like Geneformer and scGPT, we confirmed that scEMB’s performance adheres to the scaling law principle. This principle suggests that pretraining on larger and more diverse corpora consistently enhances predictive power. As scEMB was pretrained on hundreds of experimental datasets, it encountered various batch effects, technical artifacts, and individual variability during training, which ultimately improved its robustness and generalization across datasets. Larger pretraining corpora allowed scEMB to develop deeper, more predictive models capable of addressing the complexity of real-world biological data.

Finally, the introduction of *in silico* perturbation analysis opens new avenues for using scEMB in drug response prediction and cellular behavior modeling. scEMB’s ability to predict how cells will respond to various perturbations positions it as a powerful tool for future therapeutic discovery and precision medicine applications.

Moving forward, we plan to expand scEMB’s capabilities further. Leveraging generative modeling, scEMB can implicitly capture gene-gene interactions through its embeddings and attention maps, enabling the exploration of Gene Regulatory Networks (GRNs). We propose GRN inference workflows that utilize both pretrained and fine-tuned versions of scEMB, where the gene embeddings reflect dataset-level interactions, and attention maps reveal specific gene activation patterns across diverse cell states. By validating these inferred networks against established biological data, we demonstrated scEMB’s potential for gene program discovery and its ability to uncover previously unknown regulatory pathways.

## DATA AVAILABILITY

All the data generated or analyzed in this study is available from the authors upon reasonable request.

## ACKNOWLEDGEMENTS

**FUNDING**

This research was partially supported by National Institutes of Health grants awarded to Y.D R21AG087299.

